# Identification and characterization of the hypoxia response regulatory elements in the accessible genome

**DOI:** 10.1101/2020.04.24.059105

**Authors:** Xi Lu, Xingqi Chen

## Abstract

Hypoxia is commonly observed in the solid tumor and contributes to the drug resistance in cancer therapy. Deciphering the epigenetic feature under the hypoxia condition in the solid tumor is critical for us to understand the tumorigenesis and design the precision therapy. Using the time series of ATAC-seq data under the hypoxia treatment from the epithelia cells, we identified the hypoxia response regulatory elements (HRREs) in the accessible genome. We found that these different HRREs have unique genomic features and are enriched with different transcriptional factors (TFs). Our study provides insights into the chromatin structure response to the hypoxia treatment and identifies useful genomic features for a better understanding of the hypoxia biology in the solid tumor.

## Introduction

Hypoxia is commonly observed in the solid tumor^1–4^. The cancer cells in the solid tumor are found to alter their behaviors and metabolism under the hypoxia microenvironment, and further resist the cancer therapy^5,6^. In the molecular level, hypoxia induces the production of the hypoxia-inducible factors (HIFs) and other transcriptional factors (TF), modulates the chromatin structure, and further changes the gene expression in the cancer cells^7–10^. Thus, a detailed characterization of the chromatin structure changes under the hypoxia could help us to understand the tumorigenesis in the hypoxia microenvironment of the solid tumor and to design a precision therapy.

ATAC-seq technology is a powerful tool to decipher the chromatin structure with high sensitivity^11^. Previously, we demonstrated that the low level of oxygen could modulate the chromatin accessibility in the mouse breast tumor cell line 4T1 with ATAC-seq technology, and we observed that the openness of a large number of regulatory elements are dynamically changed during the time series of hypoxia treatment. We concluded that the hypoxia microenvironment alone could reshape the transcriptional factor (TF) motif openness in the genome wide^12^. However, to better understand the gene regulation under the hypoxia environment, a detailed characterization of the dynamic changes of the hypoxia response gene regulatory element is needed. Here, we performed a systematic annotation of the hypoxia response gene regulatory elements in the 4T1 cells, and identified three different types of hypoxia response gene regulatory elements (HRREs). We also found that there are different genomic features and TFs bindings on different types of HRREs.

## Results

### Identification of the hypoxia response gene regulatory elements

To characterize the hypoxia response gene regulatory elements, we included all the ATAC-seq data from the 4T1 cells under 1% O2 for various time points (0, 6, 12, 24, 36, 48, and 72 h)^12^. We used the 0h as the normoxia control for the time series comparison. When we compared the ATAC-seq peaks from each time point of hypoxia treatment with normoxia control, we identified 6,750 significant differential peaks (|logFC|>1, FDR<0.01) (**Figure 1a**, See **methods**). The openness of these significant differential peaks is modulated with the hypoxia condition; thus, these 6,750 peaks are termed hypoxia response gene regulatory elements (HRREs). Among these HRREs, a large proportion of regions, 5,015/6,750 (74.3%), start to open under the hypoxia condition, and only a small part of regions, 1,735/6,750 (25.7%), start to close with the hypoxia treatment (**Figure 1a**).

**Figure 1.**
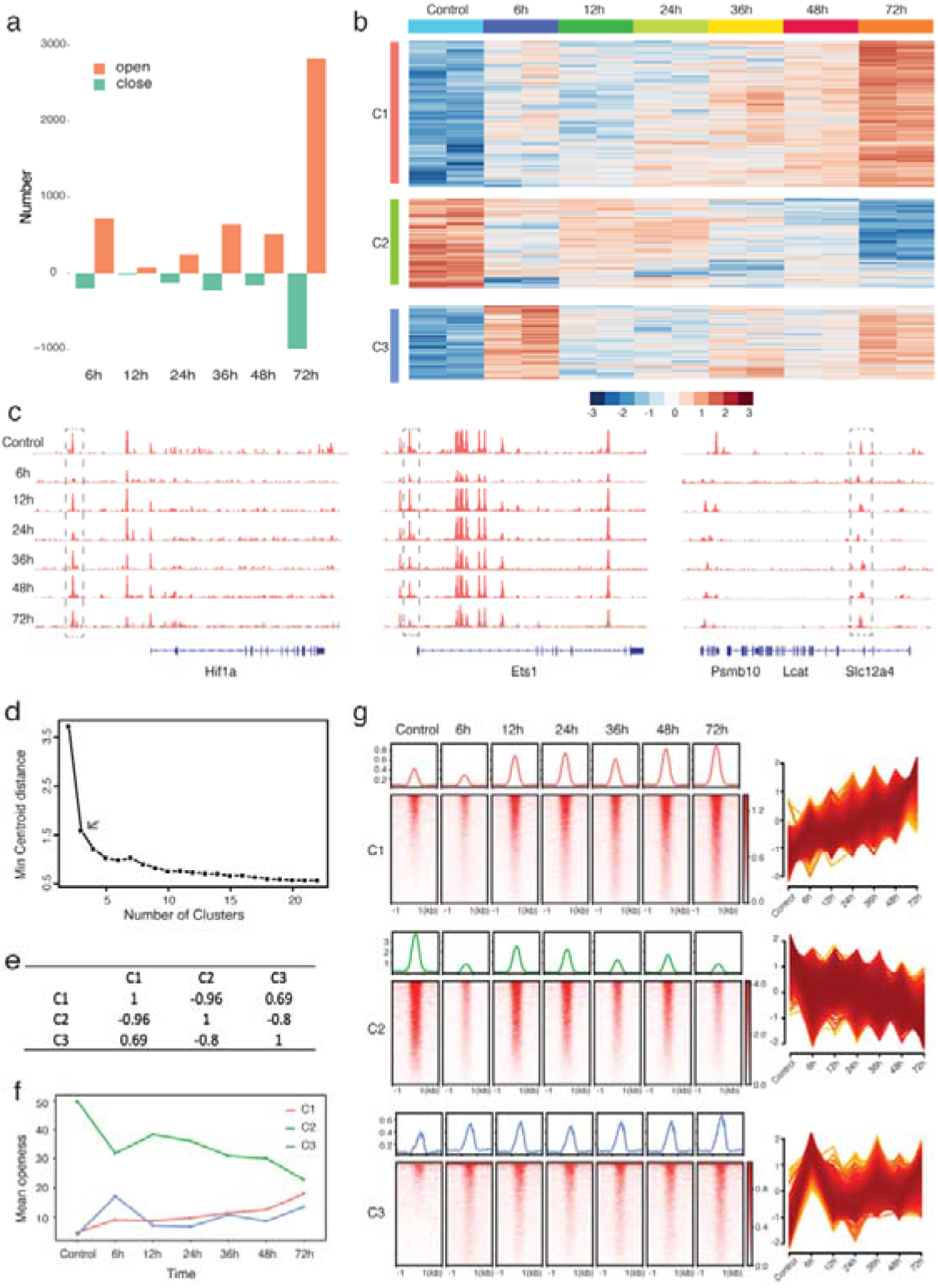
Identification of hypoxia response gene regulatory elements. a) Total accessible chromatin sites from different time points of hypoxia treatment were compared to the control (under normoxia condition). The more accessible chromatin sites are colored in orange, and the more close chromatin sites are colored in green. b) The cluster of the hypoxia response gene regulatory elements (HRREs) following the time series of hypoxia treatment. C1 = cluster 1, C2 = cluster 2, and C3 = cluster 3. c) The representative genome browser track from the different cluster of HRREs. cluster1 (Hif1a), cluster 2 (Est1), and cluster 3 (Slc12a4). The grey box indicates the location of HRREs. d) The centroid distance curve to determine the number of clusters for HRREs. e) The pearson correlation among different HRREs clusters. f) The average openness (calculated by RPKM) at each time point for each HRRE cluster. g) Left: ATAC-seq signals for cluster 1 (C1), cluster 2 (C2), and cluster 3 (C3) at different time points. Right: the chromatin openness curve at different time points.

Next, we investigated the dynamics of these HRREs across the time points (see **Methods**). Interestingly, we found the openness of 4,722 HRREs follows some trend to change during the hypoxia treatment. With the unbiased cluster methods^13^, we identified three different types of changes among these 4,772 HRREs, and named them clusters 1-3 (**Figure b-e**). The HRREs in cluster 1 (n = 2,229) came to open gradually under the hypoxia treatment compared with the control and reached the highest accessibility at 72h. We observed that the HRREs in cluster 2 (n = 1,358) closed gradually during the hypoxia treatment. We also found that the openness of HRREs in cluster 3 (n = 1,135) fluctuated during the time series of hypoxia, but had the highest accessibility after the 72h hypoxia treatment (**Figure 1f, 1g**).

### Characterization of the hypoxia response gene regulatory elements

Different types of HRREs under the hypoxia condition indicated different parts of genome responses differently to the hypoxia microenvironment. To understand the detailed mechanism of how different HRREs react differently to the hypoxia, we explored the genomic features of different HRREs in a systematic manner. When we performed the genomic annotation with all the HRRE peaks, and three different HRREs separately (**Figure 2a**), we found that the genomic annotation of HRREs in clusters 1 and 3 were similar to all the merged accessible peaks (**Figure 2a**), but the genomic annotation of HRREs in cluster 2 was quite different from the others (**Figure 2a**). In cluster 2, around 25% of the HRREs are from the promoter-TSS regions. This observation is further confirmed from the distance annotation strategy on the different groups of HRREs (**Figure 2b**), where it is clearly shown that more than 25% of the HRREs in cluster 2 are located within 1k distance from the transcriptional starting site. At the same time, the proportion of intergenic regions or distance between 10K-100k in cluster 2 is significantly less than the other clusters. Since the HRREs in the clusters start to close under the hypoxia treatment, our result indicates that a big proportion of promoter regions in cluster 2 react to the hypoxia treatment. It was reported that the histone modification, H3K27ac, is associated with a higher activation of transcription and is found at both the proximal and distal regions of the transcription start site ^14^ Thus, we assumed that the HRREs in cluster 2 are enriched with the H3K27ac histone modification. When we used the H3K27ac ChIP-Seq data from the normaixa condition to calculate the H3K27ac distribution in the HRREs from each cluster (See **Methods**) ^15^, we found indeed that there is a significant enrichment of H3K27ac in cluster 2 compared with clusters 1 and 3 (**Figure 2c**). The significant enrichment of H3K27ac in cluster 2 indicates that a big proportion of transcriptional active sites in the normoxia become silent in the hypoxia condition.

**Figure 2.**
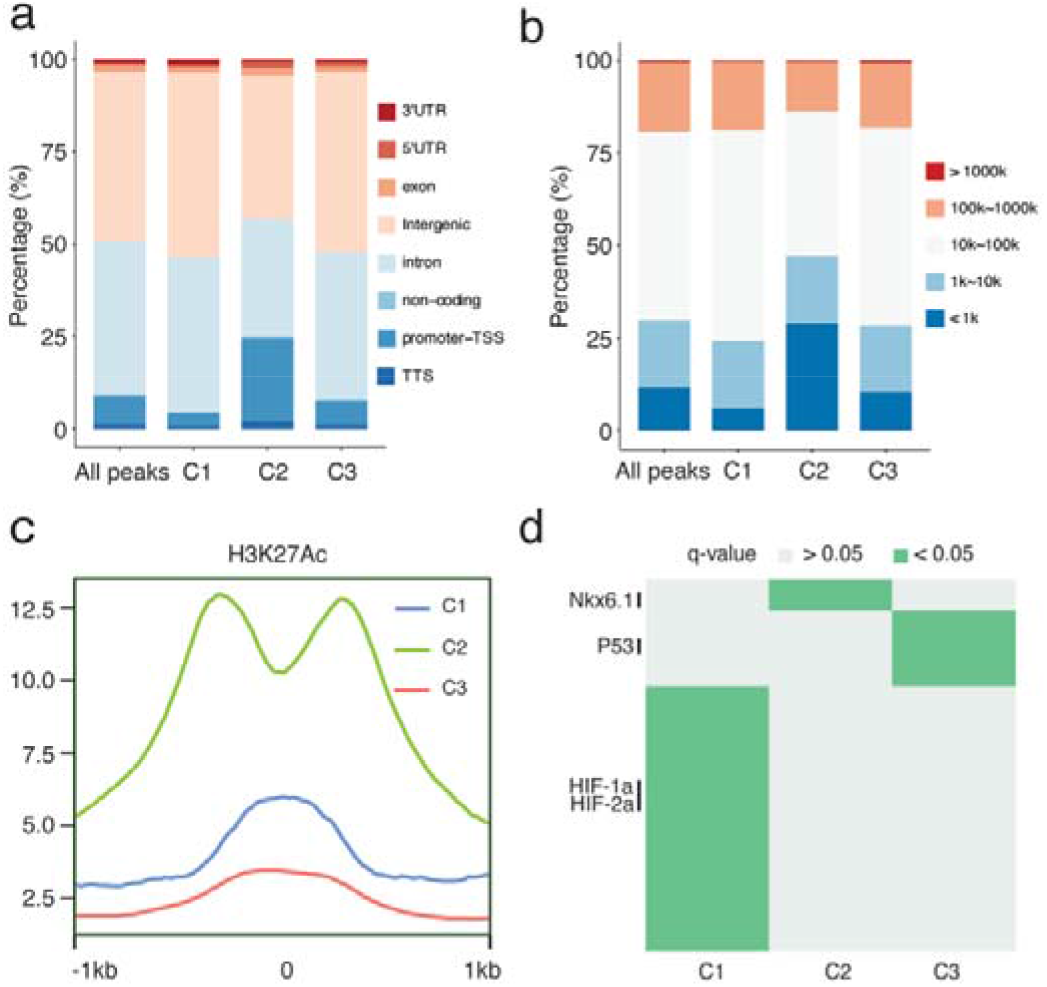
Characterization of hypoxia response gene regulatory elements. a) The genomic annotation of different hypoxia response gene regulatory elements (HRREs) clusters. C1 = cluster 1, C2 = cluster 2, C3 = cluster 3, and All = all the accessible peaks from pool samples of all time points. b) The annotation of the distance to transcription start sites (TSSs) from each HRRE clusters: cluster 1 (C1), cluster 2 (C2), and cluster 3 (C3). c) The enrichment of Histone marker H3k27ac for each HRRE cluster: 1 (C1), cluster 2 (C2), and cluster 3 (C3). d) The heatmap of enriched transcriptional factor motifs on each cluster of HRREs. C1 = cluster 1, C2 = cluster 2, and C3 = cluster 3.

It was shown that the accessibility of genomic regulatory elements is regulated by the transcription factors binding on the genome^16,17^. To understand what types of transcriptional factors regulate the different clusters of HRREs, we performed a motif enrichment for each cluster of HRREs and identified the unique transcriptional factors (TF) for each cluster (**Figure 2d**). (See **Methods**). We found that there are 152 TFs enriched in cluster 1(**Table 1**), 76 TFs enriched in cluster 2(**Table 2**), and 91 TFs enriched in cluster 3(**Table 3**) (See **Methods and Supplementary materials**). Interestingly, there are 42 shared TFs enriched in all the clusters (significant, q<0.05) (**Table 4**), and the common TFs in three different clusters demonstrated that there are common mechanism for three HRREs under the hypoxia treatment. After excluding the common TFs, we obtained a list of unique TFs identified for each cluster (See **Table 1-3**), and these specific TFs regulate the dynamic changes of chromatin accessibility in each cluster under the hypoxia treatment. Even Clusters 1 and 3 have the tendency of opening under the 72h hypoxia treatment, but the unique list for their TFs is quite different. There are 52 unique TFs for cluster 1, and 6 TFs for cluster 3. Surprisingly, the hypoxia induced protein factors, HIF1a and HIF2a, are only enriched in cluster 1, not the others (**Figure 2d**). The enrichment of hypoxia factors in cluster 1 probably explains why it is continuously open during the entire treatment time. Even cluster 3 has a tendency to stay open after 72 h; however, there is some fluctuation in openness during the time series treatment, which is probably because of the lack of hypoxia factor regulation on the HRREs in cluster 3. At the same time, it also indicates that the HRREs in cluster 3 are regulated differently from cluster 1 (**Figure 2d**). In cluster 2, 15 unique TFs were identified. The Motif bHLHE41, Nkx6.1, and CRE could be seen when regulating the chromatin structure^18–20^, and it indicates that the decrease in the chromatin accessibility in cluster 2 is regulated by these 15 TFs under the hypoxia environment.

In summary, the unique genomic feature and TFs enrichment in the different clusters of HRREs potentially reflect the different mechanisms in the reactions to the hypoxia microenvironment.

## Discussion

It was clearly shown that the hypoxia condition affects the gene expression^21^. Numerous studies have demonstrated that the hypoxia microenvironment in the solid tumor contributes to cancer development and progression^6,14,22,23^. The chromatin structure plays an important role in controlling gene expression^24^. Thus, being able to decipher the chromatin regulation under the hypoxia is vital to understand the solid tumorigenesis and to design a potential therapy strategy. Here, we performed the fine characterization of how the genomic regulatory elements respond to the hypoxia across the time series hypoxia treatment in the mouse epithelial cell line. We also identified three types of hypoxia response regulatory elements (HRREs) in the genome, and found that these different types of HRREs with different genomic features are enriched with different TFs. Our study provides insights into the chromatin structure response to the hypoxia treatment and identifies useful genomic features for a better understanding of the hypoxia biology in the solid tumor.

## Acknowledgements

This work is supported by the Swedish Research Council (VR-2016-06794, VR-2017-02074 to X.C), Beijer Foundation, Jeassons Foundation, Petrus och Augusta Hedlunds Stiftelse, Göran Gustafsson’s prize for younger researchers, Vleugel Foundation, and Uppsala University.

## Author contributions

X.L. and X.C. designed the study, and wrote the manuscript together. X.C. supervised the study.

## Materials and Methods

### Data availability

All the data used in this study were downloaded from the Gene Expression Omnibus under accession number GSE112091.

### Data processing

ATAC-seq raw data is aligned to mm9, using Bowtie2^25^ with the ‘-very-sensitive’ parameter. The properly paired reads with mapping quality larger than 30 were kept, and the duplication reads were removed with Picard. Peak calling was performed with MACS2 using the parameters –q 0.01 –nomodel –shift 0, and the peaks overlapping with the genome black list in the ENCODE were discarded.

### Identification of the differential peaks and clusters

Deseq2^26^ was used to identify the differential peaks by comparing the ATAC-seq peaks from each time point of hypoxia treatment (6h, 12h, 24h, 36h, 48h, and 72h) with ATAC-seq peaks from the normal controls. The differential peaks list is identified with the following cu-toff with: p-value < 0.01, FDR < 0.01, and the absolute value of log2 fold change > 1. The unbiased clusters from the differential peaks were tested with Mfuzz^13^. The Pearson correlation of each cluster was used to verify the independence of each cluster. The heatmaps of each cluster were generated using Deeptools^27^. Gene-element associations were then filtered to a distance of 100kb from element to TSS with HOMER^28^. KEGG and GO analyses for each cluster were conducted with GREAT^29^.

### Genomic annotation and TF enrichment

The genomic annotation of each cluster was performed based on the genomic segmentation of 3’UTR, 5’UTR, exon, intergenic, intron, non-coding, promoter-TSS, and transcription terminal sites. The distance annotation of each cluster was chosen by using the transcription start site of genes as the reference point, and the genome being separated into 5 parts: <1kbp, 1~10kbp, 10~100kbp, 100~1000kbp, and >1000kbp. The H3K27ac ChIP-seq data of 4T1 cells is downloaded from Gene Expression Omnibus^15^. The differential peaks from each cluster were annotated with H3K27ac histone marker^15^. Transcription factors for each cluster were determined using HOMER with a cut-off of q-value < 0.05. The unique TFs for each cluster are identified.

